# Robust Adaptive Deep Brain Stimulation Control of Non-Stationary Cortex-Basal Ganglia-Thalamus Network Models in Parkinson’s Disease

**DOI:** 10.1101/2023.08.20.554043

**Authors:** Hao Fang, Stephen A. Berman, Yueming Wang, Yuxiao Yang

**Affiliations:** MOE Frontier Science Center for Brain Science and Brain-Machine Integration, Zhejiang University, Hangzhou, 310000, Zhejiang, China; State Key Laboratory of Brain-machine Intelligence,Zhejiang University, Hangzhou, 310000, Zhejiang, China; College of Computer Science and Technology, Zhejiang University, Hangzhou, 310000, Zhejiang, China; Department of Neurosurgery, Second Affiliated Hospital, School of Medicine, Zhejiang University, Hangzhou, 310000, Zhejiang, China; NHC and CAMS Key Laboratory of Medical Neurobiology, Zhejiang University, Hangzhou, 310000, Zhejiang, China; Qiushi Academy for Advanced Studies, Zhejiang University, Hangzhou, 310000, Zhejiang, China; Department of Electrical and Computer Engineering, University of Central Florida, Orlando, 32816, Orlando, USA; College of Medicine, University of Central Florida, Orlando, 32816, Orlando, USA

**Author notes:** Email addresses:* (Yueming Wang), (Yuxiao Yang).

**Keywords:** Closed-loop deep brain stimulation, Parkinson’s disease, Nonlinear and non-stationary neural dynamics, Adaptive estimation, Robust control

## Abstract

Closed-loop deep brain stimulation (DBS) is a promising therapy for Parkinson’s disease (PD) that works by adjusting DBS patterns in real time from the guidance of feedback neural activity. Current closed-loop DBS mainly uses threshold-crossing on-off controllers or linear time-invariant (LTI) controllers to regulate the basal ganglia (BG) beta band oscillation power. However, the critical cortex-BG-thalamus network dynamics underlying PD are nonlinear, non-stationary, and noisy, hindering the accurate and robust control of PD neural dynamics using current closed-loop DBS methods. Here, we develop a new robust adaptive closed-loop DBS method for regulating cortex-BG-thalamus network dynamics in PD. We first build an adaptive state-space model to quantify the dynamic, nonlinear, and non-stationary neural activity. We then construct an adaptive estimator to track the nonlinearity and non-stationarity in real time. We next design a robust controller to automatically determine the DBS frequency based on the estimated PD neural state while reducing the system’s sensitivity to high-frequency noise. We adopt and tune a biophysical cortex-BG-thalamus network model as a testbed to simulate various nonlinear and non-stationary neural dynamics for evaluating DBS methods. We find that under different nonlinear and non-stationary neural dynamics, our robust adaptive DBS method achieved accurate regulation of the BG beta band oscillation power with small control error, bias, and deviation. Moreover, the accurate regulation generalizes across different therapeutic targets and consistently outperforms state-of-the-art on-off and LTI DBS methods. These results have implications for future designs of clinically-viable closed-loop DBS systems to treat PD and other neurological and neuropsychiatric disorders.

## 1. Introduction

Parkinson’s disease (PD) is one of the most common neurological disorders with an estimated global prevalence of more than 8 million [1]. PD results from the degeneration of dopaminergic neurons in the substantia nigra [2] and can lead to severe motor and cognitive symptoms [3]. It is believed that the network interactions among the cortex, the basal ganglia (BG), and the thalamus are closely related to PD motor symptoms [2, 4, 5]. Therefore, direct modulation of the cortex-BG-thalamus network activity provides a promising therapy for PD. Deep brain stimulation (DBS) is considered as the primary neuromodulation treatment method. DBS works by delivering continuous electrical stimulation pulses via electrodes implanted at focal BG targets, such as the globus pallidus internus (GPi) and the subthalamic nucleus (STN) [6, 7, 8]. Clinical results have shown that DBS can alleviate motor symptoms in PD patients, especially when the outcomes of regular drug treatment are not satisfactory [6, 8, 9].

Effective, efficient, and safe control of motor symptoms in PD patients requires precise and robust DBS of the cortex-BG-thalamus network. However, such a design of DBS is challenging for at least the three following reasons. First, the neuronal network is a dynamic structure in which the neural activities exhibit rich temporal patterns, leading to variational motor symptoms over time [4, 10]. Second, the network consists of both local and global neural interactions that make the network neural activity nonlinearly coupled [11, 12]. Third, the network activities and interactions can further be affected by time-varying and non-stationary internal and external factors such as psychiatric state variations [13, 14], circadian rhythm [15, 16], and stimulation-induced plasticity [17]. Therefore, it is difficult for standard DBS treatment of PD to achieve precise control of such a dynamic, nonlinear, and non-stationary neuronal network.

DBS treatments for PD have greatly advanced over the past few decades but have only partially addressed the above challenges. Current DBS mainly works in an open-loop manner [18, 19, 20, 21, 22, 23], i.e., a fixed pattern of stimulation is first determined by the clinician via trial- and -error and then continuously delivered regardless of symptom or neural activity variations over time. Such open-loop DBS can suffer from insufficient control of symptom dynamics, undesired side effects, and high device battery power consumption [19, 20]. Recently, closed-loop DBS strategies have been proposed as an improvement over open-loop DBS [24, 25, 26, 27, 28, 29, 30, 31, 32]. Closed-loop DBS uses real-time neural activity as feedback to guide the delivery of stimulation and holds promise to achieve more accurate and effective DBS treatment. The feedback neural activity is usually selected to be correlated with PD symptoms, such as the beta band oscillations (13–35Hz) in BG [4]. The closed-loop DBS method currently used in clinical settings works in a threshold-crossing “on-off” manner, where stimulation is triggered when the beta oscillation power exceeds a predefined threshold [24, 25, 26]. The on-off DBS addresses the dynamic changes of neural activity to some extent but lacks the ability to precisely control the neural dynamics due to its relatively simple control strategy. Linear time-invariant (LTI) controllers [27, 28, 29, 31, 32, 33] have been developed and tested in simulations as an improvement over on-off controllers. Such linear DBS methods typically use offline system identification to fit a dynamic model that quantifies the dynamic neural response to DBS. The offline fitted dynamic model is then used to design online LTI controllers, such as the proportional-integral (PI) controller [31, 32, 33] or the linear quadratic regulator (LQR) [29, 34]. Several recent simulation results have shown that LTI controllers can achieve better regulation performance of neural dynamics than on-off controllers [31, 34].

However, the linear DBS methods have limited ability to adapt to nonlinear neural dynamics [35]. Meanwhile, they use a time-invariant design, which is not robust to the non-stationarity in neural activity. Above limitations can cause linear DBS methods to have variable control performance across different treatment targets, e.g., biased control and large oscillations at some targets [31, 36].

Therefore, it is imperative to design a new DBS control method that can simultaneously address the dynamics, nonlinearity, and non-stationarity in the cortex-BG-thalamus network for PD. Recently, an adaptive PI controller has been proposed to address nonlinear neural dynamics in PD, but only tested to control limited target levels of beta oscillation power [37, 38, 39]. Also, it is not tested against non-stationary neural dynamics. Our prior work has proposed a theoretical robust adaptive neuromodulation framework based on ℒ_1_ control methods [40], in which we have focused on investigating the theoretical stability properties of the controller without testing it for specific neurological disorders.

Here, using our prior ℒ_1_ neuromodulation framework, we develop a robust adaptive DBS method for precise control of dynamic, nonlinear, and non-stationary cortex-BG-thalamus network activity in PD. Specifically, the contributions of this paper are as follows:

- We construct a realistic nonlinear and non-stationary DBS simulation testbed. We adopt and tune a well-established biophysical cortex-BG-thalamus neuronal network model [4] to simulate the ground-truth neural activity in response to DBS in PD. The original model already includes highly nonlinear neural spiking dynamics. We additionally change the neural connection strength over time to simulate more realistic non-stationary neural dynamics in PD.
- We build an adaptive state-space model to describe the cortex-BG-thalamus network dynamics. Different from prior LTI state-space models, we explicitly include nonlinear and uncertain terms to model non-linearity and non-stationarity, respectively.
- We develop a robust adaptive DBS method, which consists of a conventional linear feedback term, a new adaptive estimator, and a new robust controller. Different from prior LTI DBS methods, the adaptive estimator recursively estimates and tracks the model’s nonlinearity and non-stationarity in real time. The robust controller improves the robustness of the closed-loop system by reducing the sensitivity to high-frequency noise and disturbances.
- We comprehensively evaluate the robust adaptive DBS method using the simulation testbed. We show that the estimated adaptive state-space model tracks the nonlinear and non-stationary cortex-BG-thalamus network dynamics in PD. Also, the robust controller accurately regulates the BG beta oscillation power to the therapeutic target, resulting in significantly smaller control error, bias, and deviation than state-of-the-art on-off and linear DBS methods. Moreover, such accurate regulation can be generalized to track a wide range of therapeutic targets.
- Our results have implications for designing future clinically-viable closed-loop DBS systems for more effective treatments of PD.

The rest of the paper is organized as follows. In section 2, we describe the construction of the non-stationary cortex-BG-thalamus simulation testbed, the design of the robust adaptive DBS method, and the simulation experimental setup. Section 3 presents the results where we systematically compare the proposed robust adaptive DBS method with state-of-the-art on-off and linear DBS methods. In section 4, we discuss the contributions and limitations of our work. Section 5 concludes the paper.

## 2. Material and Methods

### 2.1 DBS simulation testbed construction using dynamic, nonlinear, and non-stationary cortex-BG-thalamus neuronal network models

The motor symptoms in PD are closely related to the cortex-BG-thalamus network [4, 5]. Thus, we adopt and modify a well-established conductance-based nonlinear biophysical model of the cortex-BG-thalamus network [4] to simulate the diseased brain under PD conditions. The original model includes the cortex (Ctx), the striatum (Str), the subthalamic nucleus (STN), the globus pallidus (GP), and the thalamus (Th). Each brain region further consists of one or two groups of neurons: Ctx contains excitatory (eCtx) and inhibitory neurons (iCtx); Str contains direct striatal projections (dStr) and indirect striatal projection (idStr); GP includes external part (GPe) and intern part (GPi); STN and Th consist of one group of neurons. Each group has 10 neurons, and the neurons are interconnected by excitatory or inhibitory connections (see figure 1A).

**Figure 1.**
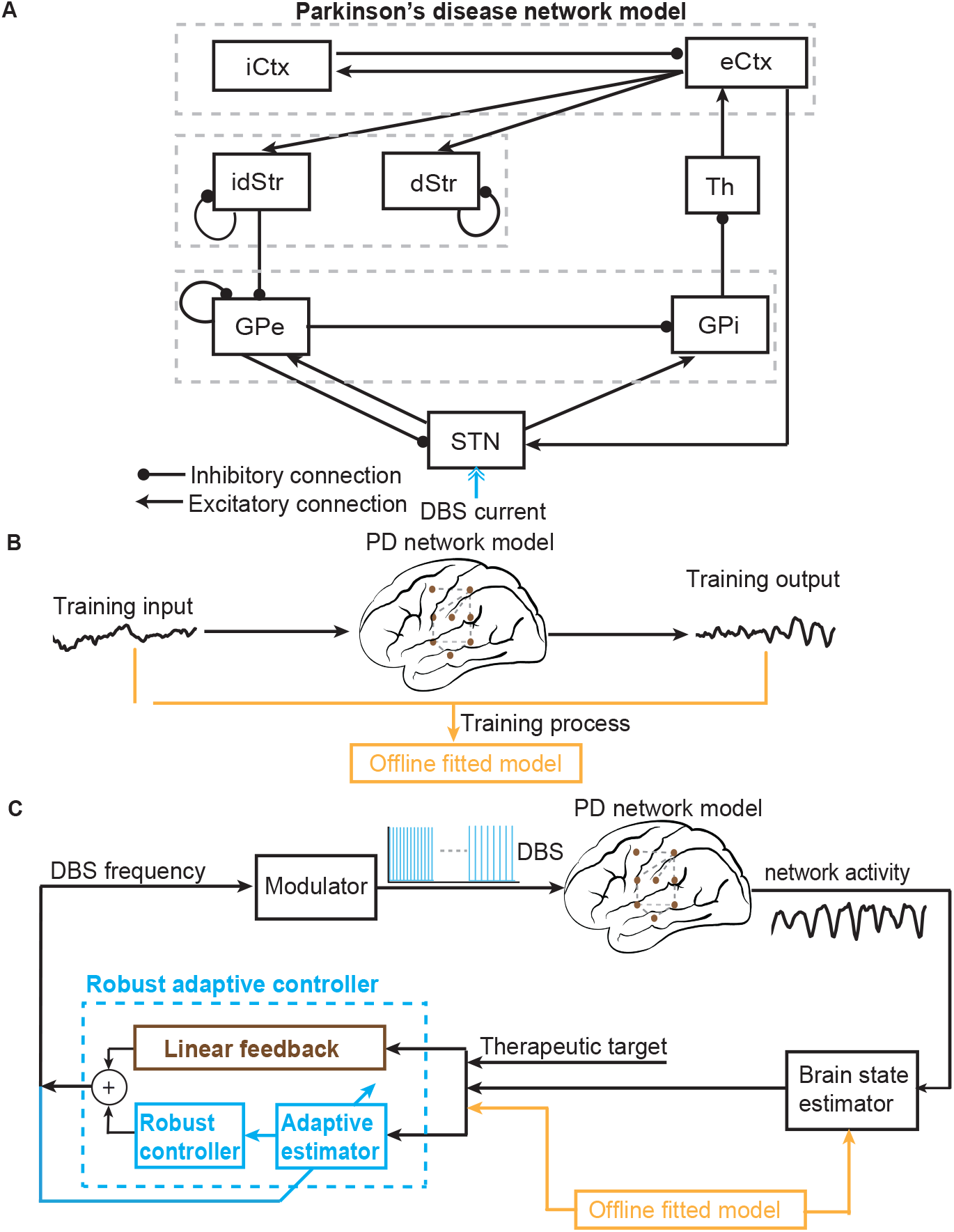
Schematic diagrams of the PD network model and the closed-loop robust adaptive neuromodulation system. (A) Illustration of the cortex-basal ganglia-thalamus network model in Parkinson’s disease. Neurons are interconnected with excitatory and inhibitory projections. (B) The training process in offline model fitting. The input is taken as the frequency of DBS and the output is the GPi beta band power. (C) The architecture of the closed-loop robust adaptive neuromodulation system. It consists of a brain state estimator, a robust adaptive controller, and a modulator. The brain state estimator estimates the PD state in real time. The robust adaptive controller consists of a linear feedback term, an adaptive estimator, and a robust controller. The robust adaptive controller takes the estimated PD state as feedback to determine the DBS frequency in real time. The modulator uses the computed DBS frequency to generate DBS pulse trains.

Specifically, the membrane potential dynamics of each neuron are modeled by conductance-based Hudgkin-Huxley-type nonlinear ordinary differential equations. For example, the following equation describes the membrane potential *v*_STN_ of a STN neuron:

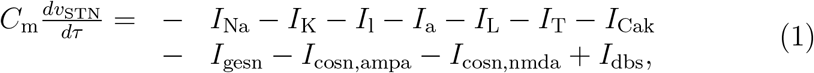

where *τ* denotes continuous time; *C*_*m*_ is the membrane capacitance; *I*_Na_, *I*_K_, *I*_l_ are sodium ionic currents, potassium ionic currents, and leakage currents; *I*_a_ is an A-type potassium ionic current; *I*_L_ is a L-type calcium current; *I*_T_ is a T-type calcium current; *I*_Cak_ is a calcium-dependent potassium current that is dependent upon the intracellular calcium concentration; *I*_gesn_ is the synaptic input current from GPe neurons; *I*_cosn,ampa_ and *I*_cosn,nmda_ are the Ctx-STN synaptic currents mediated by AMPA receptor and NMDA receptor, respectively; *I*_dbs_ is the external DBS input current. Each current uses a conductance-based dynamic model. For example, the inhibitory synaptic current *I*_gesn_ is modeled by

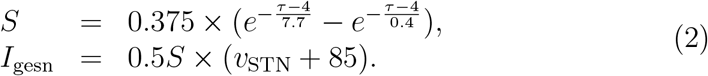

We see that the exponential operation in the above synaptic current example introduces nonlinear temporal dynamics in the differential equation. Therefore, the entire model consists of much more complicated nonlinear neural dynamics. Readers who are interested can refer to the detailed full model [4]^1^.

To simulate a pathological Parkinsonian state in the above cortex-BG-thalamus model, one can tune a key variable pd that changes the neural coupling strength between several key regions in the model:

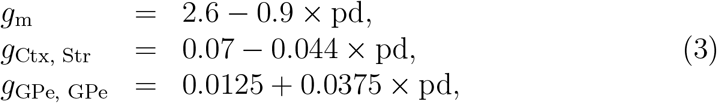

where *g*_m_ is the M-type potassium conductance among Str neurons; *g*_Ctx, Str_ is the cortical-striatal coupling strength; *g*_GPe, GPe_ is the internal GPe coupling strength. The healthy (normal) state is simulated by setting pd = 0, and a typical severe PD state is simulated by setting pd = 1. We note that the above cortex-BG-thalamus network model has been tested against clinical experimental data and shown to be useful in describing diseased neural dynamics underlying PD by many previous studies [4, 31, 41].

The original cortex-BG-thalamus network model [4] assumes that the key variable pd in (3) is fixed at a constant value of 1 for the Parkinsonian state, leading to a time-invariant and stationary model. Such a stationary model serves as a simulation testbed that has been widely used to evaluate various DBS methods for PD [4, 31, 41, 37]. However, a fixed pd cannot characterize the non-stationary PD neural dynamics that have been observed in practice, such as the changes in neural connectivity due to the change in disease condition [17], circadian rhythm [16], psychiatric state variations [13], and stimulation-induced plasticity [12]. Therefore, we make critical modifications to the original cortex-BG-thalamus network model to simulate the above biophysically-plausible non-stationary neural dynamics. Specifically,

1. We simulate a scenario where the disease condition makes non-stationary changes over time by switching pd = 1 to pd = 0.5 over time because the reduction of PD symptom severity may happen over time;
2. We simulate the non-stationarity caused by circadian rhythm or psychiatric state variations via setting pd as a periodic sinusoidal function of time, i.e., pd = sin(*ωt* + *θ*), where *t* is the time step, *ω* is the angular frequency and *θ* is the phase.
3. We simulate the non-stationarity caused by stimulation-induced plasticity via setting pd as a nonlinear function of the real-time DBS input to the model, i.e., pd = *f* (*u*_*t*_), where *f* (·) is a nonlinear function and *u*_*t*_ represents the real-time DBS input, which can itself be time-varying as in the closed-loop DBS case.

The detailed simulation setups are included in section 2.5. With the above modifications, we extend the original cortex-BG-thalamus network model to include various types of biophysically plausible non-stationary neural dynamics. Subsequently, we use the modified nonlinear and non-stationary cortex-BG-thalamus network models as the final simulation testbeds for evaluating different DBS methods.

### 2.2 Problem formulation of closed-loop DBS control of cortex-BG-thalamus network models

We first formally define the input and output variables of the cortex-BG-thalamus network model in the context of closed-loop DBS control. Prior studies have shown that the oscillatory neural activities, especially the beta band oscillation, in BG are closely related to PD symptoms [4, 42]. Particularly, GPi neurons can exhibit exaggerated beta band oscillations under the Parkinsonian state. Therefore, following prior studies [4, 42], we choose GPi beta band power as the key signal needed to be regulated by DBS and define the GPi beta band power as the output of the cortex-BG-thalamus network model. Specifically, we compute the output *y*_*t*_ as the average beta band power of the voltage potential across all GPi neurons:

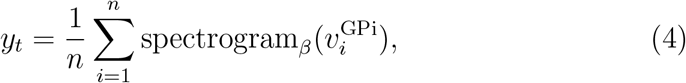

where 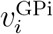 is the continuous voltage potential of the *i*th GPi neuron, *n* = 10 is the total number of GPi neurons. We use the standard multitaper spectral analysis [43] (represented by the function spectrogram_*β*_ (·)) to compute the beta band power of 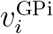 where we take the discrete time step *t* to be 0.1s, the sliding window to be 1s, the time-bandwidth product to be 3, and the number of tapers to be 5.

For the input of the model, prior studies have shown that the pulse train frequency is one of the most important DBS parameters that affect DBS treatment outcome [4, 31, 29]. Therefore, following prior studies [4], we take the input to the cortex-BG-thalamus network model as the frequency of the DBS pulse train,

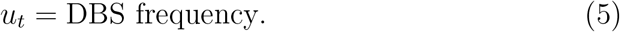

To be consistent with the discrete time step of the output (*t* = 0.1*s*), we enable the DBS frequency to change every *t* = 0.1s. Next, the corresponding continuous DBS pulse train *I*_dbs_ at STN can be constructed by a pulse train with a fixed amplitude of 300*µ*A/cm^2^ and a fixed duration of 0.1 s, and a time-varying pulse frequency of *u*_*t*_.

With the definition of input (5) and output (4) of the cortex-BG-thalamus network model, our goal is to design a closed-loop DBS method that can determine the input DBS frequency *u*_*t*_ in real time to regulate the output GPi beta band power *y*_*t*_ to track a pre-defined constant therapeutic target value *y*^*∗*^. In practice, the target value *y*^*∗*^ can be selected by the clinician based on experience [24, 26]. In prior simulation studies, DBS methods are usually evaluated against a single target value of *y*^*∗*^. Here, we evaluate DBS methods in tracking different values of *y*^*∗*^ at section 3.3.

### 2.3 Current closed-loop DBS methods

#### 2.3.1 On-off DBS

The state-of-the-art closed-loop DBS that have been tested in PD patients typically uses an “on-off” control strategy [24, 25, 26]. In on-off DBS, a neural feature correlated with PD symptoms such as the BG beta band power is first chosen as the feedback signal. Then a threshold value of the feedback signal is chosen ad-hoc. In real-time closed-loop DBS control, a fixed pattern of DBS is triggered whenever the feedback signal crosses the threshold. In our case, we can choose the feedback signal as the model output *y*_*t*_ and the threshold as the therapeutic target *y*^*∗*^. Therefore, on-off DBS works by setting

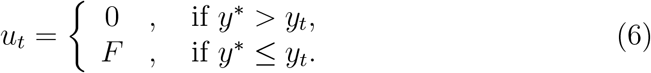

Here *F* is the fixed stimulation frequency, which is typically chosen as the high frequency of 130 Hz accordingly to prior clinical studies [12, 25, 44]. As an improvement of traditional open-loop DBS, on-off DBS has been shown to reduce side-effects and battery power consumption [24, 25]. However, the on-off DBS method does not model the neural dynamics in response to stimulation, thus the simple on-off control strategy has limited abilities to address nonlinear and non-stationary neural dynamics in PD [12, 25].

#### 2.3.2 Linear DBS

To further improve on-off DBS, several closed-loop DBS methods based on LTI modeling of neural dynamics have been proposed for PD and tested in simulations [27, 28, 29, 31]. The core idea of these methods is that a mathematical model that quantifies the neural response to stimulation can provide useful information to construct linear feedback controllers. Specifically, these methods first use an offline model fitting experiment to construct a LTI dynamic model to quantify the neural response to DBS. In the context of PD, the offline model fitting experiment delivers a specific open-loop input *u*_*t*_ and simultaneously records the corresponding output *y*_*t*_ in response to the input. Then, the input-output dataset is used to fit an LTI dynamic model (see figure 1B). Various types of LTI dynamic models have been used such as the auto-regressive model [27, 31] and the state-space model [29]. Next, the LTI dynamic model is used to design various types of LTI DBS controllers such as the Proportional-integral (PI) controller [31] and the linear quadratic regulator (LQR) [29]. Finally, in a separate online control experiment, the LTI controllers are tested in regulating the output neural activity *y*_*t*_ toward the therapeutic target *y*^*∗*^ .

Here, based on our prior work [29], we present a typical linear DBS framework that fits a LTI state-space model and enable design a PI controller or LQR. The LTI state-space model is

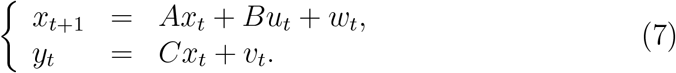

Here, 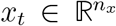 represents the Parkinsonian state [29]. *u*_*t*_ *∈* ℝ is the frequency of DBS. 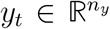 is the GPi beta band power spectral. *w*_*t*_ and *v*_*t*_ are white Gaussian noise that represents modelling error. *w*_*t*_ and *v*_*t*_ have zero mean and a joint covariance matrix 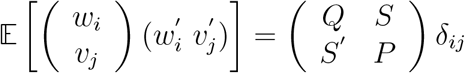 with *δ*_*ij*_ = 1 if *i* = *j* and 0 otherwise, 𝔼 [·] and _*′*_ denoting the expectation and the matrix transpose operator, respectively. The model parameters are the *A, B, C, Q, S, P* matrices and are fitted by the aforementioned input-output dataset (see figure 1B and [29]).

Using the offline fitted model (7), one can design a linear PI controller

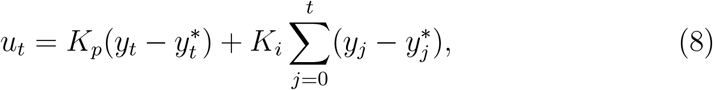

where *K*_*p*_ and *K*_*i*_ are the proportional and integral gains, respectively. *K*_*p*_ and *K*_*i*_ can be either set by the user in an ad-hoc manner [31] or optimized by simulating the offline fitted model (7) and choosing the option that leads to the best-simulated control performance [40].

Another typical LTI controller takes the following state feedback form

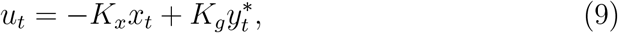

where *K*_*x*_ and *K*_*g*_ are the feedback and feedforward gains, respectively. Both gains can be computed as functions of the fitted model parameters in (7).

For example, we use the well-known LQR solution for *K*_*x*_ and *K*_*g*_,

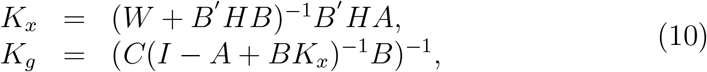

where *W* is a user-defined positive definite matrix, *H* is solved from the algebraic Riccati equation *H* = *A*′ *HA* + *C*′ *C − A*′ *HB*(*W* + *B*′ *HB*)^*−*1^*B*′ *HA* [45]. The Parkinsonian state *x*_*t*_ can be estimated by constructing a brain state estimator such as a static linear estimator [40]

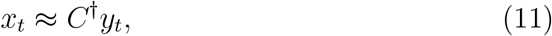

where *C*^*†*^ = (*CC*^*T*^)^*−*1^*C* is the pseudo-inverse, or a more sophisticated dynamic Kalman estimator [46, 47, 48].

Simulation studies have shown that linear DBS improves over the on-off DBS for PD [31, 29, 27]. However, linear DBS uses linear models and controllers, which are not sufficient to capture and address the complex nonlinear dynamics in PD [31, 28, 29, 40]. Moreover, the linear models and controllers are time-invariant, meaning their parameters are fixed in the online control experiment. However, non-stationary neural dynamics can happen in real time and thus cannot be captured by time-invariant models or controllers. This can lead to biased control with large oscillations [31, 40, 49, 47]. Therefore, a new closed-loop DBS method is needed to address the nonlinear and non-stationary neural dynamics in PD.

### 2.4 Robust adaptive DBS

To overcome the aforementioned limitations of on-off and linear DBS, we develop a robust adaptive DBS method based on the ℒ_1_ neuromodulation framework proposed by our prior work [40]. Briefly, we first build an adaptive state-space model and explicitly include nonlinear and uncertain terms to model nonlinearity and non-stationarity, respectively. We then develop an adaptive estimator to recursively estimate and track the model’s nonlinearity and non-stationarity in real time. Finally, we develop a robust controller that compensates for the model’s nonlinearity and non-stationarity while reducing control sensitivity to noise. The overall robust adaptive control architecture is taken as the sum of a linear feedback term and a robust adaptive augmentation term,

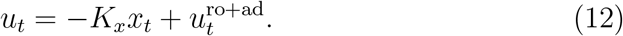

Next, we briefly present the adaptive state-space model, the adaptive estimator, and the robust controller. The readers can refer to our prior work [40] for the detailed mathematical theories underlying the design.

#### 2.4.1 Adaptive state-space model

We extend the LTI state-space model in (7) to be the following adaptive state-space model

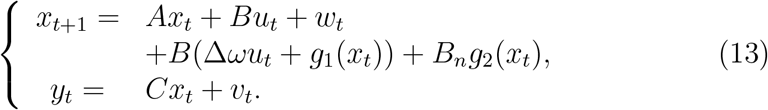

Note that the time-invariant parameters *A, B, C, Q, S, P* are obtained in the same way as in the LTI state-space model in (7) from offline model fitting. *B*(∆*ωu*_*t*_ + *g*_1_(*x*_*t*_)) and *B*_*n*_*g*_2_(*x*_*t*_) are the new and critical terms added in the state equation that represent nonlinear and non-stationary model uncertainty that happen in real time and cannot be captured by the offline fitted model (7) prior to stimulation. Therefore, the entire model is adaptive. The first term *B*(∆*ωu*_*t*_ + *g*_1_(*x*_*t*_)) characterizes the uncertainty lying in the range space of the input matrix *B*, ∆*ω* is an unknown input gain uncertainty, *g*_1_(·) is an unknown nonlinear function; the second term *B*_*n*_*g*_2_(*x*_*t*_) describes the uncertainty lying in the null space of control, i.e., 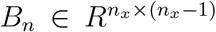 satisfies *B*′ *B*_*n*_ = **0** (**0** represents a zero vector of corresponding dimension) and rank([*B, B*_*n*_]) = *n*_*x*_, *g*_2_(·) is an unknown nonlinear function. The nonlinear and uncertain terms ∆*ω, g*_1_(*x*_*t*_), *g*_2_(*x*_*t*_) can be estimated in real time by the following adaptive estimator.

#### 2.4.2 Adaptive estimator

We design a parametric adaptive estimator to track nonlinear and uncertain terms ∆*ω, g*_1_(*x*_*t*_), *g*_2_(*x*_*t*_) in (13). First, we parametrize *g*_1_(*x*_*t*_) and *g*_2_(*x*_*t*_) by using a time-varying Taylor approximation

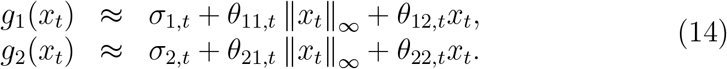

Here, 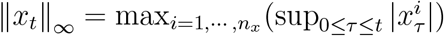 is the truncated ℒ_*∞*_ norm [40] (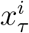 represents the *i*th component in *x*_*τ*_), which represents the maximum value of the state variable across all its components (the outer max operation) and across all time steps from the initial time step 0 to the current time step *t* (the inner sup operation). The time-varying parameters after parametrization 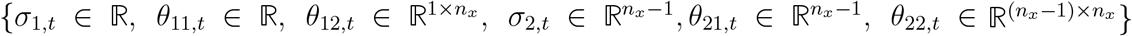 need to be estimated. Thus, we develop an adaptive estimation algorithm. Specifically, we plug the above approximations (14) and control architecture (12) into the state equation (13), and simplify it to get

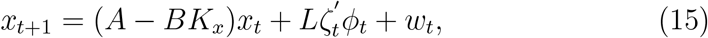

where *L* = [*B, B*_*n*_], *ζ*_*t*_ = [*ζ*_1,*t*_, *ζ*_2,*t*_], *ζ*_1,*t*_ = [*ω*_*t*_, *σ*_1,*t*_, *θ*_11,*t*_, *θ*_12,*t*_]′, *ζ*_2,*t*_ = [**0**, *σ*_2,*t*_, *θ*_21,*t*_, *θ*_22,*t*_]′ and 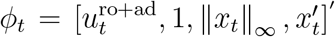 Here, *ζ*_*t*_ contains all the time-varying parameters that need to be estimated. We then rewrite (15) as

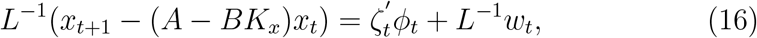

which is a linear regression model of the parameters *ζ*_*t*_ with *ϕ*_*t*_ as the regressor. We apply the standard recursive least squares method [40] to compute the estimate of *ζ*_*t*+1_ as

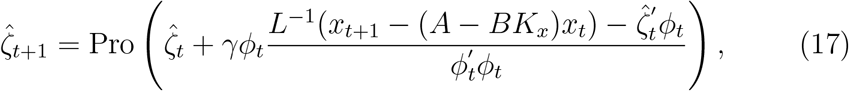

where 0 *< γ≤* 1 is the adaptation rate and Pro(*·*) is the standard projection operator [40] to ensure the estimations are all bounded. Finally, we can estimate range space uncertainty ĝ_1,*t*_ = ∆*ωu*_*t*_ + *g*_1_(*x*_*t*_) and null space uncertainty ĝ_2_(*x*_*t*_) as

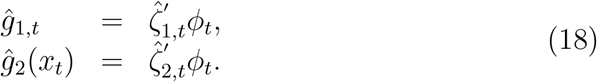

#### 2.4.3 Robust controller

Based on the control architecture (12) and the adaptive estimates of model uncertainty (18), we can derive the explicit robust adaptive controller as the following:

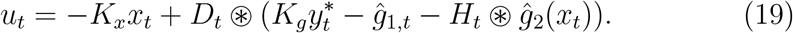

where ⊛ is the linear convolution operator, *D*_*t*_ and *H*_*t*_ are the impulse responses of two linear filters that enforce robust control as detailed below. Due to the limited space, we omit the detailed derivations and only provide some key considerations. Interested readers can refer to our prior work [40] for the details of theoretical derivations.

First, the final purpose of the controller is to use state feedback to regulate the neural activity to track the therapeutic target *y*^*∗*^, thus we inherit the state feedback term *−K*_*x*_*x*_*t*_ and feedforward term *K*_*g*_*y*^*∗*^ from the LTI controller (9).

Second, the controller needs to address the model’s nonlinearity and uncertainty. This is done by including the term *−*ĝ_1,*t*_ to cancel the range space nonlinearity and uncertainty, and the term *−H*_*t*_ ⊛ ĝ_2_(*x*_*t*_) to cancel the null space nonlinearity and uncertainty. Note that ĝ_2_(*x*_*t*_) is in the null space of the input matrix and cannot be completely canceled by the controller. Therefore, the controller only aims to cancel ĝ_2_(*x*_*t*_) at steady state, and this is realized by the linear filter *H*_*t*_. To this end, the *z* transform of *H*_*t*_ can be derived as *H*_*z*_ = (*C*(*zI − A* + *BK*_*x*_)^*−*1^*B*_*n*_)(*C*(*zI − A* + *BK*_*x*_)^*−*1^*B*)^*−*1^.

Third, the adaptive estimates ĝ_1,*t*_ and ĝ_2_(*x*_*t*_) contain estimation errors that are caused by high-frequency noise and disturbances. Therefore, any cancellation of model uncertainty should only happen within the bandwidth of the closed-loop system since aggressive cancellation outside the bandwidth is essentially trying to cancel irreducible noise, and can lead to instability [40]. Given that brain networks generally behave similarly to a low-pass system [50, 10], we use a low-pass filter *D*_*t*_ to reduce the high-frequency errors in ĝ_1,*t*_ and ĝ_2_(*x*_*t*_). The *z* transform of *D*_*t*_ takes as a first-order low-pass filter 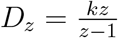 with *k* being the filter gain. With the key low-pass filter *D*_*t*_, we have rigorously proved in our prior work [40] that the controller (19) is stable and robust to high-frequency noise and disturbances.

To this end, we combine the linear feedback term, adaptive estimator, and robust controller to form our robust adaptive DBS method. We summarize the robust adaptive DBS method in Algorithm 1 (also see figure 1C).

### 2.5 Simulation setups

Based on the cortex-BG-thalamus network simulation testbed that we build in section 2.1, we conduct comprehensive Monte Carlo simulations to evaluate the proposed robust adaptive DBS (19) and compare with the on-off DBS (6), the linear PI DBS (8), and the LQR DBS (9).

#### 2.5.1 Offline part

In the offline model fitting experiment prior to closed-loop DBS control, we assume that the patient is under severe PD condition and set the key parameter pd = 1 [4]. We then generate an open-loop input pattern by drawing the DBS frequency *u*_*t*_ randomly from a uniform distribution on [0, 200] Hz at each time step. The total random input time step *T* is 3000.

##### Algorithm 1 The robust adaptive DBS algorithm

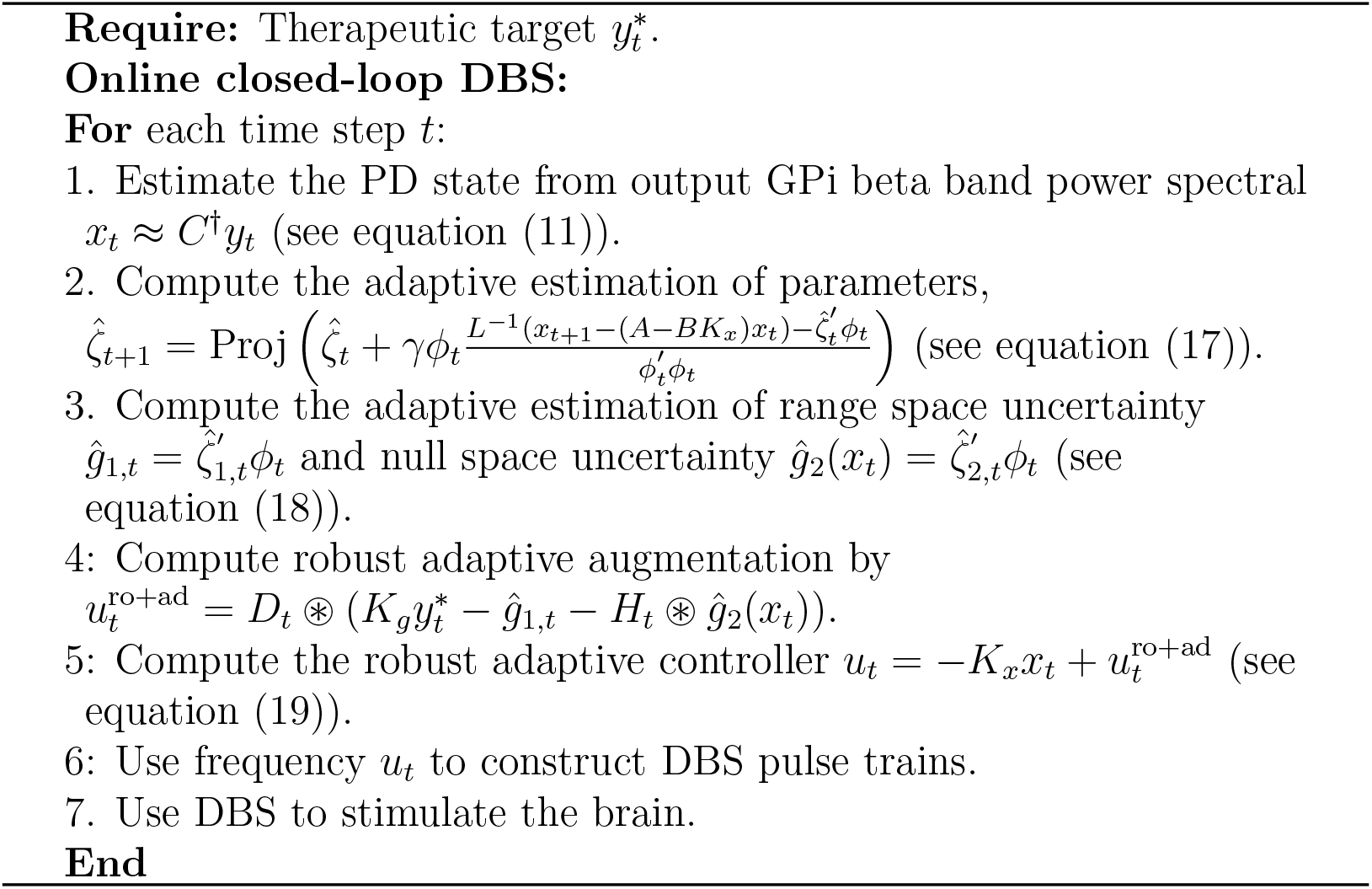

We next apply the random input pattern to the cortex-BG-thalamus network model to sufficiently excite the neural dynamics and collect the responding output GPi beta band power *y*_*t*_. We organize the input-output dataset as (*u*_1_, *y*_1_), (*u*_2_, *y*_2_), …, (*u*_*T*_, *y*_*T*_) and use it to fit a LTI state-space model (7) using the standard subspace identification method [51].

#### 2.5.2 Online part

For online closed-loop DBS, we run the following simulations for each closed-loop DBS method.

- First, we consider a stationary case and assume that the disease condition of the patient stays the same across time, i.e., we set pd = 1 in the cortex-BG-thalamus network model throughout the online closed-loop DBS control experiment. We focus on evaluating different DBS methods in regulating the output GPi beta band power to track a constant therapeutic target *y*^*∗*^ = 150. For each DBS method, we simulate 50 independent trials with different initial conditions of the cortex-BG-thalamus network model. Each trial lasts for 15 seconds where the closed-loop DBS is not turned on until the 3rd second.
- Next, we use more realistic non-stationary cortex-BG-thalamus network models as simulation testbeds. We simulate three scenarios as detailed in section 2.1: (1) to simulate a non-stationary reduction of symptom severity over time, we switch the disease condition from pd = 1 to pd = 0.5; (2) to simulate non-stationarity caused by circadian rhythm or psychiatric state variations, we set pd = 0.5 + 0.5sin(*t* + *ϕ*) to be sinusoidal functions of time with three different phase shifts 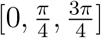for the three conductance terms in (3), respectively. Note that each pd has a mean value of 0.5 and the overall oscillation is still within the range [0, 1]; (3) to simulate non-stationarity caused by stimulation-induced plasticity, we set pd = 0.5 +0.5sin(*u*_*t*_ +*ϕ*) as a nonlinear function of the DBS input *u*_*t*_, where we again use three different phase shifts 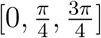 for the three conductance terms in (3), respectively. In the above three scenarios, we keep focusing on tracking a constant therapeutic target *y*^*∗*^ = 150 and simulate 50 independent trials for each DBS method.

Finally, to test if the DBS method can generalize to track different therapeutic targets, we repeat the above experiments but change the constant therapeutic target to different values. In total, we simulate 5 targets *y*^*∗*^ from {110, 130, 150, 170, 190}. For each target value and each DBS method, we also simulate 50 independent trials.

### 2.6 Performance metrics and statistical testing

To evaluate the closed-loop DBS control performance, we define the control error (CE) at time step *t* as

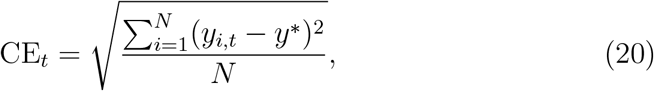

where *y*_*i,t*_ is the output signal in trial *i* and *N* = 50 is the total number of trials. We focus on the CEs at the steady state where we exclude the first 5s of CEs immediately after DBS starts to avoid the transition period. For different closed-loop DBS methods, we compare the steady-state CEs by the Wilcoxon signed-rank test. We then investigate two important factors that can contribute to CE, i.e., control bias and control deviation. We define control bias (CB) as the absolute difference between the trial-averaged output signal 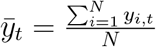 and the target *y*\*.

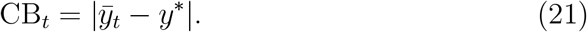

We define control deviation (CD) as the oscillation around the trial-averaged output signal, i.e., across-trial standard deviation

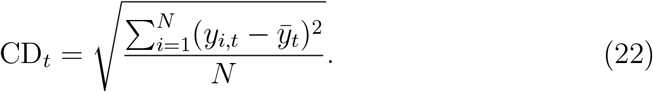

Similarly, for different closed-loop DBS methods, we compare the steady-state CEs and CDs by the Wilcoxon signed-rank test.

For the non-stationary cortex-BG-thalamus network models, we also compare how the offline LTI state-space model in equation (7) and online adaptive state-space model in equation (13) can predict the non-stationary output neural activity *y*_*t*_. In a single trial, we apply a 6s random input pattern to the non-stationary cortex-BG-thalamus network model and record the responding output neural activity *y*_*t*_. For the prediction using the offline LTI state-space model, we plug the same 6-second input into equation (7) and set all noise terms *w*_*t*_ and *v*_*t*_ to be identically zero over time and recursively compute the output as the model prediction *ŷ*_*t*_. For prediction using the online adaptive state-space model, we plug the same 6s input into equation (13), setting all noise terms *w*_*t*_ and *v*_*t*_ to be identically zero over time, and estimating *g*_1,*t*_ and *g*_2_(*x*_*t*_) with the adaptive estimator (18) over time. We then recursively compute the output as the model prediction *ŷ*_*t*_. We define the prediction error as:

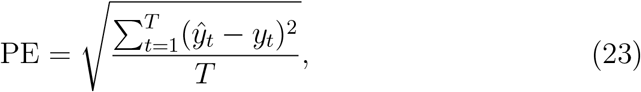

where *T* = 60 is the total time steps. We compare the offline LTI state-space model prediction and the online adaptive state-space model prediction across 50 independent trials using the Wilcoxon signed-rank test.

## 3. Results

### 3.1 Robust adaptive DBS tracks a constant therapeutic target in stationary cortex-BG-thalamus networks

We started with stationary cortex-BG-thalamus networks and qualitatively examined the control performance of the robust adaptive DBS and other closed-loop DBS methods at three different scales: the tracking of the average GPi beta band power, the change in the detailed GPi beta spectrogram, and the change in the spiking patterns of GPi neurons (the three columns in figure 2). For the robust adaptive DBS (see figure 2A), the average GPi beta power stayed high before DBS started, representing the Parkinsonian neural state. Once robust adaptive DBS turned on, the average GPi beta power started to drop and after a short transition period, it stayed close to the therapeutic target. Accordingly, in the detailed GPi beta spectrogram, we saw that under robust adaptive DBS, there was an overall power reduction across the entire beta band. For GPi neurons, before DBS turned on, the neurons fired synchronously with a beta rhythm as was typical in the Parkinsonian neural state. Robust adaptive DBS disrupted the synchronous firing pattern, leading to the reduction of GPi beta power.

**Figure 2.**
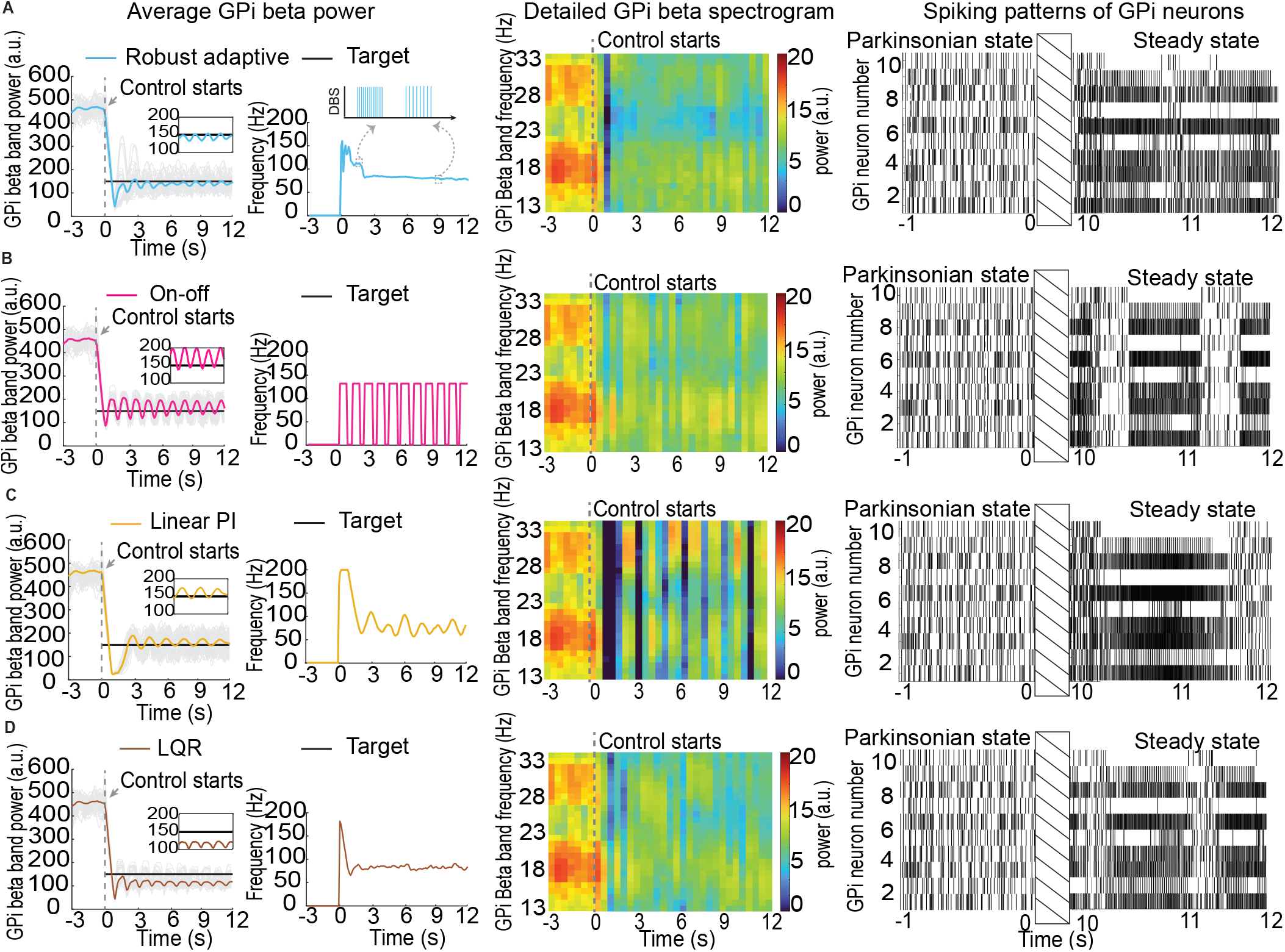
Robust adaptive DBS tracks a constant therapeutic target in stationary cortex-BG-thalamus networks. (A) Control performance of the robust adaptive DBS method. There are three columns in each row from left to right. The first column shows the controlled average GPi band beta power. Each simulation trial is in gray and the trial-averaged is in cyan. The dashed line represents the starting time point of closed-loop DBS. We zoom into the steady state as the insertion. We also show the associated control input (the controlled DBS frequency). The inserted small panel shows the actual DBS pulse train with the indicated DBS frequency. The second column shows the detailed GPi beta spectrogram. The dashed line represents the starting time point of closed-loop DBS. The third column shows the spiking patterns of GPi neurons, where each vertical line represents a spiking event and the shaded area separates the spiking patterns of the Parkinsonian state before DBS and the steady state after DBS. (B) Same with (A) but for on-off DBS. (C) Same with (A) but for linear PI DBS. (D) Same with (A) but for LQR DBS.

By contrast, the on-off DBS, linear PI DBS, and LQR performed worse than the robust adaptive DBS. For the on-off DBS (see figure 2B), the controlled average GPi beta power had a large bias and deviation around the therapeutic target. Accordingly, the detailed GPi beta spectrogram showed less reduction of power (especially at 23-28 Hz) and obvious variations over time (especially at 18-23 Hz). Moreover, on-off DBS did not sufficiently disrupt the GPi neurons’ synchronous firing pattern, especially from 10.2s to 10.5s and from 11.3s to 11.7s. For linear PI DBS (figure 2C), the controlled average GPi beta power had a small bias but the deviation stayed large. Accordingly, the detailed GPi beta spectrogram showed undesired large variations across the entire beta band and the GPi neurons’ synchronous firing pattern was not sufficiently disrupted (e.g., from 11.7s to 12s). For LQR DBS (figure 2D), the controlled average GPi beta power had a small deviation but the bias stayed large. Accordingly, the detailed GPi beta spectrogram showed overly-reduced power at 28-33 Hz and the GPi neurons’ synchronous firing pattern was again not sufficiently disrupted (e.g., from 11.1s to 11.4s).

Next, wentitatively compared the control performance of the above closed-loop DBS methods. In terms of the overall control error (figure 3 left panel), the on-off DBS had the largest control error (CE: 43.63 [41.40, 45.86], mean and 95% confidence interval) and the linear PI and LQR DBS significantly reduced the control error (Linear PI CE: 42.12 [41.19, 43.04]; LQR CE: 41.01 [39.90, 42.13], Wilcoxon signed-rank test *P <* 10^*−*10^ in both cases), confirming the results in prior studies. However, the linear PI and LQR DBS only reduced around 3.74% of the control error compared with the on-off DBS. By contrast, our robust adaptive DBS significantly reduced the control error by 29.41% compared with the on-off DBS (Robust adaptive CE: 30.80 [29.57, 32.03] vs. On-off CE: 43.63 [41.40, 45.86], Wilcoxon signed-rank test *P <* 10^*−*10^), and the control error was also significantly smaller than the linear PI DBS and the LQR DBS (Wilcoxon signed-rank test *P <* 10^*−*10^ for both comparisons).

**Figure 3.**
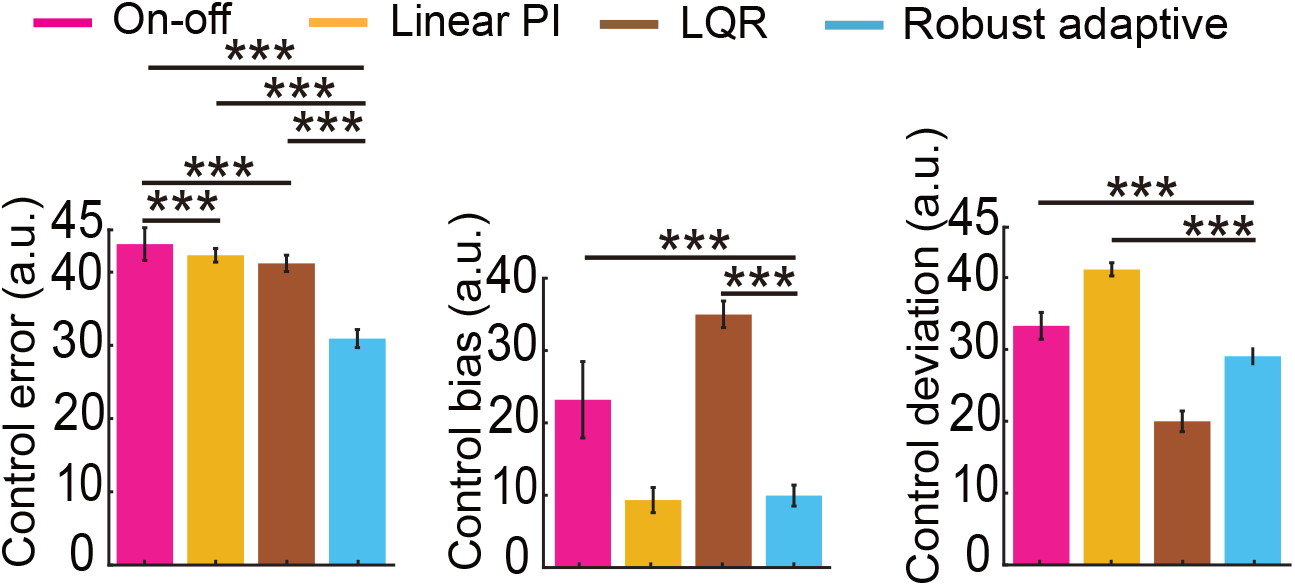
Comparison of control error, bias, and deviation for different closed-loop DBS methods for stationary cortex-BG-thalamus networks. The bar represents the mean value of each performance metric and the whiskers represent its 95% confidence interval. *∗ ∗ ∗* represents *P <* 0.0005.

We further decomposed the overall control error into control bias and control deviation (figure 3 middle and right panels). The on-off DBS had much larger control bias and control deviation than our robust adaptive DBS (Wilcoxon signed-rank test *P <* 10^*−*10^ for both comparisons). The linear PI DBS had comparable control bias with our robust adaptive DBS but its control deviation was much larger (Wilcoxon signed-rank test *P <* 10^*−*10^), leading to the overall larger control error. On the other hand, the LQR DBS had a slightly smaller control deviation than our robust adaptive DBS but a much larger control bias (Wilcoxon signed-rank test *P <* 10^*−*10^), leading to the overall larger control error. The above quantitative results confirmed the qualitative results shown in figure 2.

To summarize, for stationary cortex-BG-thalamus networks, we confirmed the previous findings that linear PI and LQR DBS methods outperformed on-off DBS. More importantly, we further demonstrated that in stationary networks, our robust adaptive DBS already significantly improved over linear PI and LQR DBS methods in tracking a constant therapeutic target.

### 3.2 Robust adaptive DBS tracks a constant therapeutic target in various types of non-stationary cortex-BG-thalamus networks

We have shown that in stationary cortex-BG-thalamus networks, our robust adaptive DBS method outperformed prior closed-loop DBS methods. Next, we turned to the more practical and challenging cases of controlling non-stationary cortex-BG-thalamus networks. We started with the first type of non-stationary cortex-BG-thalamus networks where the neural coupling strength switched as step functions over time. Specifically, we set the disease condition to be pd = 1 before closed-loop DBS and switched it to be pd = 0.5 during online closed-loop DBS (see figure 4A).

**Figure 4.**
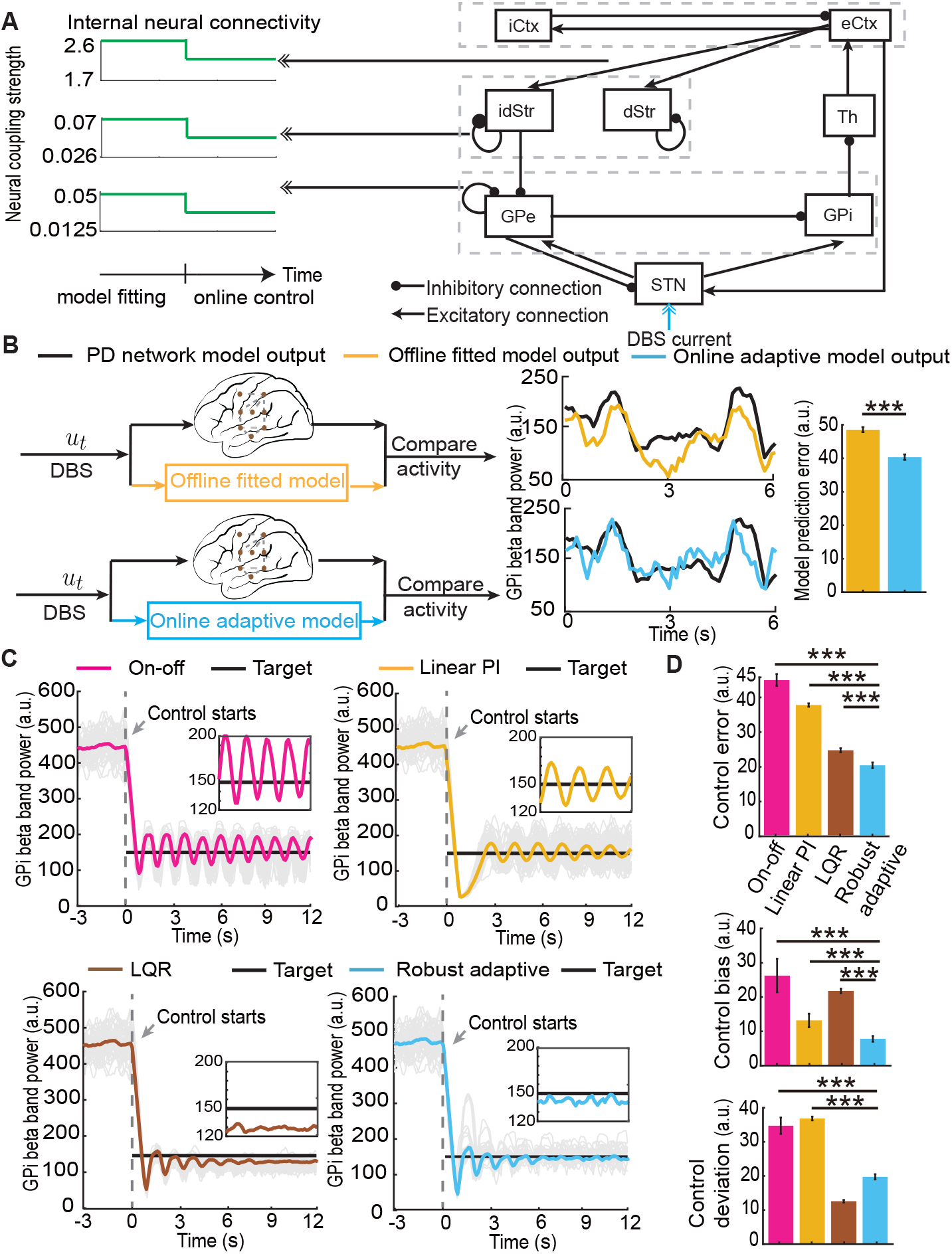
Robust adaptive DBS tracks a constant therapeutic target in the first type of non-stationary cortex-BG-thalamus networks with switching changes in the neural coupling strength. (A) The neural coupling strength switch as step functions over time. (B) Comparisons of model prediction error. The bar represents the mean value of the model prediction error and the whiskers represent its 95% confidence interval. *∗ ∗ ∗* represents *P <* 0.0005. (C) The controlled GPi beta power of different closed-loop DBS methods. The figure captions are the same as in figure 2. (D) Comparisons of control error, bias, and deviation for different closed-loop DBS methods. The figure captions are the same as in figure 3.

We first examined how well the offline fitted LTI state-space model and online adaptive state-space model predicted the non-stationary neural activity. In this case, the LTI state-space model was fitted prior to online control under the condition pd = 1. As a result, the LTI state-space model did not predict the GPi beta power well during online closed-loop control, when the disease condition was switched to pd = 0.5 (see figure 4B top panel). By contrast, our online adaptive state-space model addressed neural non-stationarity and achieved better prediction of the GPi beta power during online closed-loop control (see figure 4B bottom panel); specifically, its model prediction error was significantly smaller than the LTI state-space model (PE: 38.78 [37.63, 39.92] vs. 47.77 [46.72, 48.81], Wilcoxon signed-rank test *P <* 10^*−*10^). This result confirmed that the simulated neural activity was indeed non-stationary and our adaptive state-space model successfully tracked the non-stationarity.

Next, we turned to evaluate different closed-loop DBS methods in tracking a single constant therapeutic target (figure 4C and 4D). The linear PI and LQR DBS methods were derived from the LTI state-space model and our robust adaptive DBS method was derived from the adaptive state-space model. A more precise computational model of the cortex-BG-thalamus network should lead to a better closed-loop DBS method. Given the better prediction performance of the online adaptive state-space model, we expected that our robust adaptive DBS method should outperform the linear PI and LQR DBS methods as well as the on-off DBS method (which did not even use a model to design the controller). Indeed, we found that the on-off DBS did not track the target and had large control error, bias, and deviation (CE: 44.80 [43.08, 46.53], CB: 25.93 [21.06, 30.80], CD: 33.07 [30.77, 35.36]). The linear PI DBS reduced the control bias (CB: 12.87 [10.91, 14.85]) but the control deviation was still large (CD: 35.09 [34.57, 35.62]), thus only leading to a slightly reduced control error (CE: 37.58 [37.09, 38.08]). The LQR DBS reduced the control deviation (CD: 11.95 [11.57, 12.30]) but the control bias was still large (CB: 21.44 [20.78, 22.09]), again only leading to a slightly reduced control error (CE: 24.52 [23.93, 25.12]). By contrast, our robust adaptive DBS significantly reduced both the control bias (CB: 7.53 [6.69, 8.37]) and deviation (CD: 18.76 [18.02, 19.49]), collectively leading to a significantly smaller control error than all the other closed-loop DBS methods (CE: 20.18 [19.36, 21.00], Wilcoxon signed-rank test *P <* 10^*−*10^ for all comparisons).

We next extended the above evaluations to the second type of non-stationary cortex-BG-thalamus networks where we varied the neural coupling strength continuously over time as sinusoidal functions (figure 5A). This scenario simulated neural non-stationarity caused by circadian rhythms or psychiatric state variations. Similarly, we found that our online adaptive model had significantly smaller prediction error than the offline fitted LTI model (figure 5B, PE: 38.44 [37.42, 39.47] vs. 43.04 [42.11, 43.98], Wilcoxon signed-rank test *P <* 10^*−*10^). Consequently, our robust adaptive DBS achieved significantly smaller control error than the other closed-loop DBS methods (Figure 5C and D, CE: 24.04 [22.52, 25.57] vs. On-off CE: 35.96 [34.42, 37.50], Linear PI CE: 33.58 [31.65, 35.51], LQR CE: 30.81 [27.74, 33.88], Wilcoxon signed-rank test *P <* 10^*−*10^ in all comparisons).

**Figure 5.**
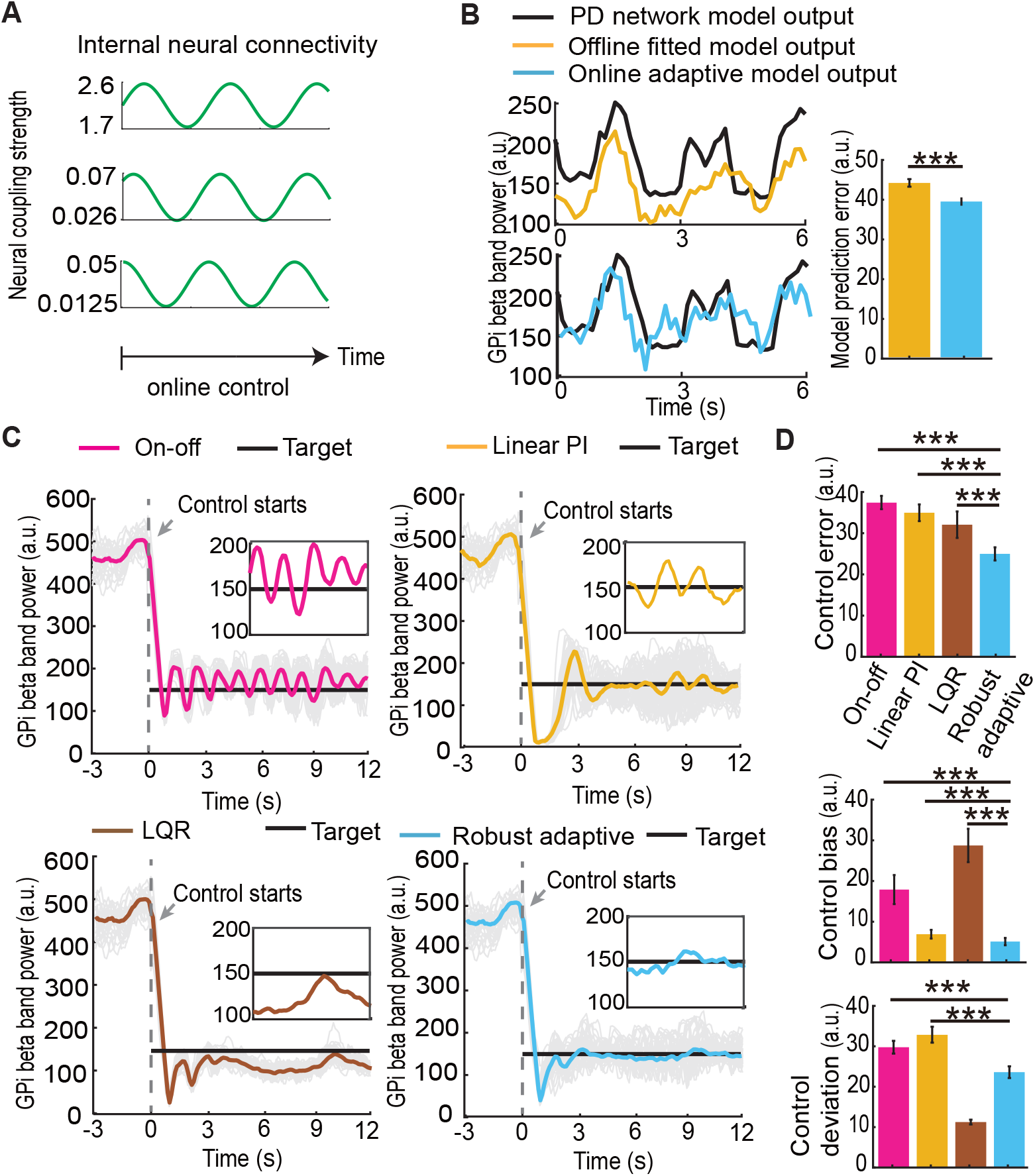
Robust adaptive DBS tracks a constant therapeutic target in the second type of non-stationary cortex-BG-thalamus networks with sinusoidal changes in the neural coupling strength. (A) The neural coupling strengths switch as sinusoidal functions over time. (B-D) The figure captions are the same as in figure 4.

Finally, we extended the evaluations to the third type of non-stationary cortex-BG-thalamus networks where we varied the neural coupling strength continuously over time as functions of the input DBS (figure 6A). This scenario simulated neural non-stationarity caused by stimulation-induced plasticity. Similarly, we found that the online adaptive model had significantly smaller prediction error than the offline fitted LTI model (figure 6B, PE: 37.61 [36.55, 38.67] vs. 42.90 [41.87, 43.93], Wilcoxon signed-rank test *P <* 10^*−*10^). Consequently, our robust adaptive DBS achieved significantly smaller control error than the other closed-loop DBS methods (figure 6C and D, CE: 28.14 [27.54, 28.74] vs. On-off CE: 30.11 [28.97, 31.25], Linear PI CE: 52.16 [51.21, 53.12], LQR CE: 33.30 [32.62, 33.99], Wilcoxon signed-rank test *P <* 10^*−*10^ in all comparisons).

**Figure 6.**
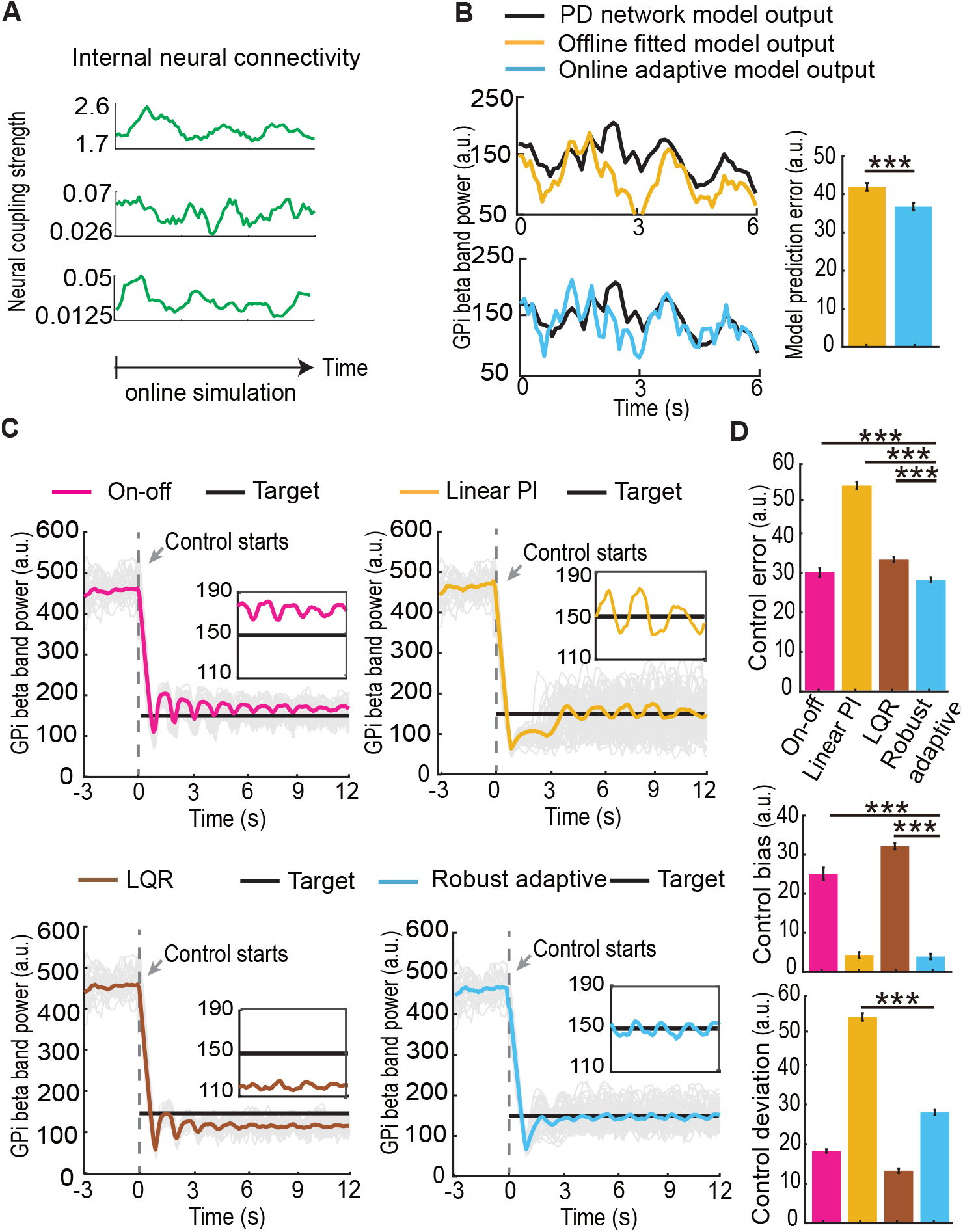
Robust adaptive DBS tracks a constant therapeutic target in the third type of non-stationary cortex-BG-thalamus networks with stimulation-induced plasticity in the neural coupling strength. (A) The neural coupling strengths change as functions of the real-time DBS frequency over time. (B-D) The figure captions are the same as in Figure 4.

In summary, for three different types of non-stationary cortex-BG-thalamus networks, we consistently showed that our online adaptive nonlinear state-space model successfully tracked the non-stationary neural activity. As a consequence, our robust adaptive DBS methods accurately tracked the ther-apeutic target and significantly outperformed the on-off DBS, the Linear PI DBS, and the LQR DBS methods.

### 3.3 Robust adaptive DBS generalizes accurate tracking across multiple therapeutic targets

We have so far evaluated closed-loop DBS methods in tracking a constant therapeutic target of 150 for the GPi beta power. In practice, different values of constant therapeutic targets should be chosen depending on different clinical requirements. Therefore, we repeated all the above simulations of controlling non-stationary cortex-BG-thalamus networks with 5 different targets {110, 130, 150, 170, 190} (figure 7). We found that the control error of the on-off DBS stayed large regardless of the target value. The linear PI and LQR DBS had smaller control errors for small target values but quickly approached the large control error (similar to the on-off DBS) as the target value increased. By contrast, the control error of our robust adaptive DBS stayed small and only slightly increased as the target value increased. Across the 5 tested target values, the control error of our robust adaptive DBS was significantly smaller than the other closed-loop DBS methods (CE: 28.09 [27.77, 28.40] vs. On-off CE: 40.88 [40.61, 41.15], Linear PI CE: 37.53 [36.95, 38.11], LQR CE: 35.73 [35.18, 36.28], Wilcoxon signed-rank test *P <* 10^*−*10^ for all comparisons). To summarize, our robust adaptive DBS accurately regulated the GPi beta power at multiple therapeutic targets and consistently outperformed state-of-the-art closed-loop DBS methods.

**Figure 7.**
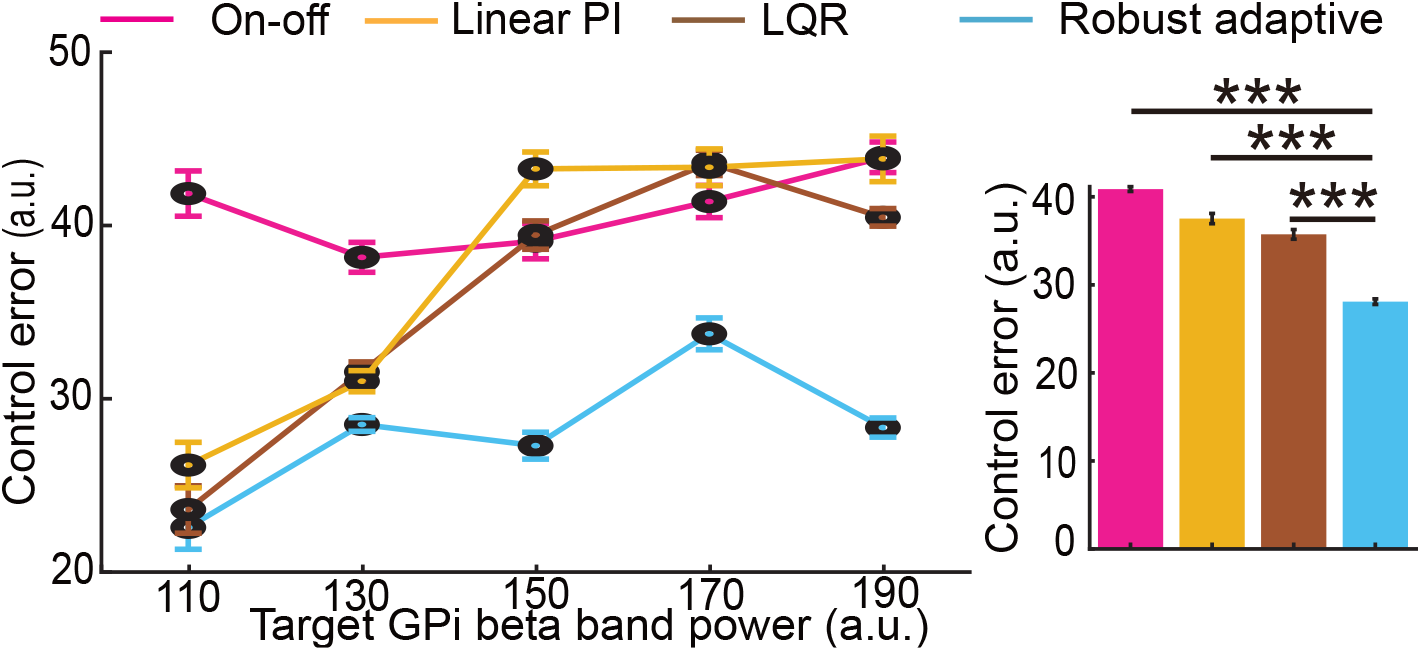
Robust adaptive DBS generalizes accurate tracking across multiple therapeutic targets. Left panel: control error as a function of a therapeutic target for different closed-loop DBS methods. The circle represents the mean value in controlling all three types of non-stationary cortex-BG-thalamus networks and whiskers represent its 95% confidence interval. Right panel: the aggregated control error across all tested therapeutic targets. Figure captions are the same as in Figure 3.

## 4. Discussion

With the long-term goal of achieving precise DBS control of motor symptoms in PD, we develop a robust adaptive closed-loop DBS method for regulating dynamic, nonlinear, and non-stationary neural activity and evaluate it using biophysical cortex-BG-thalamus network models. Our method focuses on addressing two critical aspects of neural dynamics in PD: nonlinearity and non-stationarity. Many factors can contribute to the nonlinear and non-stationary neural dynamics in PD. First, nonlinear neural dynamics naturally arise from the internal complex neuronal interactions in the cortex-BG-thalamus loop [4, 2, 28, 52]. Second, PD is a long-term neurological disorder where symptoms can evolve from mild tremors to severe postural instability as PD progresses [2, 13]. On the other hand, PD symptoms may be alleviated during DBS and/or drug treatments [7, 6]. Therefore, symptom variations can induce non-stationarity. Third, the symptom and neural dynamics of PD can also be affected by internal biological processes and rhythms such as psychiatric state [13, 14], blood pressure [53], circadian rhythm [16]. Fourth, stimulation-induced plasticity during DBS can elicit non-stationary discharge patterns in motor and sensory cortices [12]. Moreover, external factors such as body movements [54] and recording noise [55] can introduce real-time noise and disturbance during DBS. Therefore, we specifically changed the neuronal coupling strength over time in a well-established cortex-BG-thalamus network model [4] to simulate various non-stationary neural activities. Prior closed-loop DBS methods for PD have used simple “on-off” or LTI control strategies [24, 25, 26, 27, 28, 31, 32]. Our simulation results show that these prior methods have limited ability to address the nonlinear and non-stationary neural dynamics, resulting in large control bias and/or control deviation. By contrast, our robust adaptive DBS method realized accurate and robust control of various types of simulated nonlinear and non-stationary cortex-BG-thalamus neural dynamics and consistently outperformed state-of-the-art closed-loop DBS methods.

Prior developments of closed-loop DBS methods for PD have largely been evaluated in simulations that aim to regulate the neural activity to a single constant pre-defined target value [27, 28, 34, 37, 33]. Previous studies have shown that decreased beta band oscillations in BG correlate with the alleviation of PD symptoms [42, 56], but the exact therapeutic target value of beta power remains unclear and can be different in different patients [2]. Moreover, patients have heterogeneous symptoms as PD evolves over time [2]. Therefore, tracking a single target value may not be sufficient for treating PD and different target values may be needed depending on the application scenario. Our results show that the proposed robust adaptive DBS method maintained accurate tracking for several typical target values in a relatively large therapeutic regime and again consistently outperformed state-of-the-art closed-loop DBS methods across the different target values.

As a prerequisite for in-vivo animal and patient experiments, evaluating closed-loop DBS methods for PD using a biophysical cortex-BG-thalamus network model is a common practice [4, 31, 28, 41, 37]. We conducted comprehensive simulations and successfully evaluated our robust adaptive DBS method. Our next step is to move towards in-vivo testing. However, in-vivo animal and patient testing of closed-loop DBS for PD is still challenging. First, while BG beta power correlates with PD symptoms [42, 56, 44] and is frequently used as the feedback signal in closed-loop DBS [4, 31], other neural features can also indicate PD symptom severity such as the STN *γ* power [25] and the GPi firing patterns [28]. Thus, incorporating various neural features to provide more effective feedback signals is an important future direction. Second, in terms of the DBS input, clinical studies have shown that the PD treatment effect can depend on various DBS parameters such as pulse frequency, pulse amplitude, and pulse width [29, 27]. Our future work will extend the robust adaptive closed-loop method to simultaneously control these different DBS parameters. Third, we have built the robust adaptive DBS upon the adaptive state-space model. Our current adaptive estimator has assumed that the linear matrix parameters are estimated in a separate offline model fitting experiment and only estimated the other non-linear uncertain model parameters online. Our prior work has shown that such offline model fitting is feasible in animal experiments [10] but can still introduce treatment interruption [57, 49, 58]. Extending our adaptive estimator to also estimate the linear matrix parameters online and completely remove the offline model fitting experiment is another direction that requires further investigation [35, 59].

Finally, our robust adaptive DBS method provides a general design framework to develop closed-loop therapies for other neurological and neuropsychiatric disorders. Recently, there are rich explorations of effective feed-back signals and closed-loop DBS treatments for epilepsy [60], major depression [47, 61], and chronic pain [62]. Our robust adaptive DBS framework is flexible to model different types of input DBS parameters and output neural activity. Thus, it is feasible to tailor our framework to address the specific feedback signals and DBS parameters for the DBS treatment of different neurological and neuropsychiatric disorders, which is also an important future research direction.

## 5. Conclusions

Taken together, we developed a new robust adaptive DBS method for closed-loop neuromodulation treatment of PD. We conducted comprehensive simulations to evaluate its performance using dynamic, nonlinear, and non-stationary cortex-basal ganglia-thalamus neuronal network models. We found that the proposed robust adaptive DBS method accurately regulated nonlinear and non-stationary neural activity across multiple therapeutic targets, significantly outperforming state-of-the-art closed-loop DBS methods. Our results have implications for future clinically-viable closed-loop neuro-modulation system designs to treat PD and other neurological and neuropsy-chiatric disorders.

### CRediT authorship contribution statement

Yuxiao Yang: Conceptualization, Methodology, Supervision, Writing-original draft, Writing–review and editing. Hao Fang: Methodology, Conducting simulations, Analysing Data, Visualization, Writing–original draft, Writing-review and editing. Stephen A. Berman: Writing-review and editing. Yueming Wang: Writing-review and editing.

## Declaration of Competing Interest

The authors declare that they have no known competing financial interests or personal relationships that could have appeared to influence the work reported in this paper.

## Data Availability Statement

Data will be made available on request.

## Acknowledgement

This work was partly supported by grants from the National Key R&D Program of China (2018YFA0701400), Key R&D Program of Zhejiang (no. 2022C03011), the Chuanqi Research and Development Center of Zhejiang University, the Starry Night Science Fund of Zhejiang University Shanghai Institute for Advanced Study (SN-ZJU-SIAS-002), the Fundamental Research Funds for the Central Universities, the UCF COM and CECS 2021 Pilot Awards Program.

## Footnotes

1 The code for simulating the original cortex-BG-thalamus neuronal network model has been made public at http://senselab.med.yale.edu/modeldb.

## Notes

### Competing Interest Statement

The authors have declared no competing interest.

